# Clade-Specific MPXV PCR Assays

**DOI:** 10.1101/2023.04.24.538151

**Authors:** Daniel Antonio Negrón, Nicholas Tolli, Stephanie Guertin, Suzanne Wollen-Roberts, Shane Mitchell, Jared Haas, Catherine Pratt, Katharine Jennings, Bradley Abramson, Lauren Brinkac, David Ashford

## Abstract

In an evolving infectious disease outbreak, there are two priorities: rapid and accurate detection of the causative agent and characterization of its spread. The polymerase chain reaction (PCR) assay is an effective and portable diagnostic method that can quickly provide information associated with virulence, transmissibility, and pathogenicity. Compared to genomic sequencing, PCR requires less infrastructure, funding, and training. However, the development of sensitive and specific primer sequences is costly, particularly those with subspecies resolution. The recent *mpox (monkeypox) virus* outbreak underscores the need for the rapid development of clade-specific primers, particularly when there are differences in morbidity and mortality rates. Current mpox assays use primer sequences that also bind to the broader *Orthopoxvirus* genus, resulting in suspect diagnoses, delays in treatment, and poor allocation of scarce healthcare resources. Additionally, these orthopox-based primer sets cannot distinguish between different mpox clades and cannot illuminate intra-clade evolution over the course of the outbreak. Here, we present the *in silico* design and *in vitro* testing of novel clade-specific mpox assays.

**Importance:** There is an ongoing global outbreak of Mpox disease, an illness with a characteristic blistering rash progression. The Mpox virus clusters into two clades with differing levels of virulence and mortality rates. Thus, proper sample identification is critical for surveillance and the public health response programs that rely on such data. Accordingly, the US government specifically lists Clade I under the Federal Select Agent Program. Current diagnostics may fail to identify the virus or its clade membership as the genome mutates. In this work, we demonstrate an end-to-end workflow to quickly evaluate existing PCR assays and design new ones for clade-based identification with respect to large sequence databases in service of preventing a catastrophic fog-of-war from forming during an outbreak.

## Introduction

*Mpox virus*, also known as *monkeypox virus* (MPXV), is an enveloped, linear, double-stranded DNA virus in the *Orthopoxvirus* genus and *Poxviridae* family (1). To reduce geographical stigma, the former Congo Basin (Central African) clade is now referred to as Clade I, and the former West African clade as Clade II (2). There are two subclades within Clade II, IIa and IIb, with the latter corresponding to variants circulating during the 2022 global outbreak (2). Clade II was originally thought to be less severe with milder disease, less deaths, and limited human-to-human transmission compared to Clade I (3, 4).

The exact *mpox virus* reservoir is unknown, although multiple species are carriers, including rodents and primates (5, 6). Transmission to humans is typically related to direct contact with infected animals, body fluids, or respiratory droplets, with some evidence supporting aerosol and fomite transmission (7). Mpox was first identified in humans in 1970 in the Democratic Republic of the Congo (DRC) two years after the eradication of smallpox (8). The cessation of vaccinia vaccination, waning immunity following smallpox eradication, and increased human migration have increased the risk of mpox and catalyzed the recent global outbreak (9). As of February, 1, 2023, the United States has reported 30,123 cases and 28 associated deaths, with 85,777 total global cases since May 2022 (10). Historically, fatality rates were 10% for Clade 1 cases and 1% for Clade II cases, however Clade IIb cases have experienced a lower fatality rate of 0.1%. This difference may be due to several factors, such as healthcare access, divergence, and selection bias (11).

Mpox disease in humans manifests as an initial febrile prodrome 3-17 days after exposure, with headache and fatigue, potentially accompanied by enlarged lymph nodes (10). At rash onset, the fever often declines and lesions of varying distribution and number develop through the macular, papular, vesicular, and pustular phases until scabbing for a period of 14-28 days (10, 12). In recent years, atypical presentations have included only genital lesions and absence of fever (10). The key clinical characteristics of mpox disease present similarly to smallpox disease and other illnesses associated with vesicular lesions, thereby complicating diagnosis (12). Improved molecular diagnostics are critical to rapid, correct diagnoses, targeted treatments, and controlling spread of the disease.

*Orthopoxvirus* infections are difficult to characterize by serology due to cross-reactivity between other genus members, making molecular testing by polymerase chain reaction (PCR) the preferred diagnostic method for mpox, given its accuracy and sensitivity (12). Mpox also differs from other viral pathogens in that standard specimens needed for accurate diagnosis are skin lesions, dry crusts, or biopsy, instead of the more typical and accessible blood, serum, and sputum samples, meaning that PCR-based diagnostics are only effective when samples are taken during the active rash phase of the illness (12).

The recent influx of mpox cases and the difficulty of diagnosing atypical infections outside of endemic regions highlights the need for rapid identification to assist with diagnosis and case management to decrease further community spread (6, 13). Molecular dating, phylogenetic, and coding region analysis can assist with understanding potential transmission chains and epidemiological factors (14, 15). Furthermore, sequencing data assists medical countermeasure development, suitability, and effectiveness (16). However, the current laboratory infrastructures where mpox is endemic need improvement for effective surveillance, treatment, and prevention of the disease, including portable molecular testing capabilities (6, 9, 12). Here we describe novel primer sets to identify mpox and discern between Clade I and Clade II variants in the field.

## Results

The protocol identified 55 generic (pan-clade), 690 Clade I, and 1,595 Clade II MPXV assay candidates from 60,702 pan-clade, 116,124 Clade I, and 161,989 Clade II BioVelocity-identified signature sequences respectively. Table S1 lists the complete set of control and candidate assay sequences and Table S2 and Table S3 list the corresponding alignment statistics and global-local alignments from the PSET evaluation. Figure 1 displays the amplicon and primer ranges of the tested PCR assays with reference genome context and variation and Table 1 lists the corresponding primer and probe sequences. Figure S1 similarly displays the ranges of all assays. Of the tested assays, no indels were observed to occur within primer regions, only very few SNPs.

**Figure 1.**
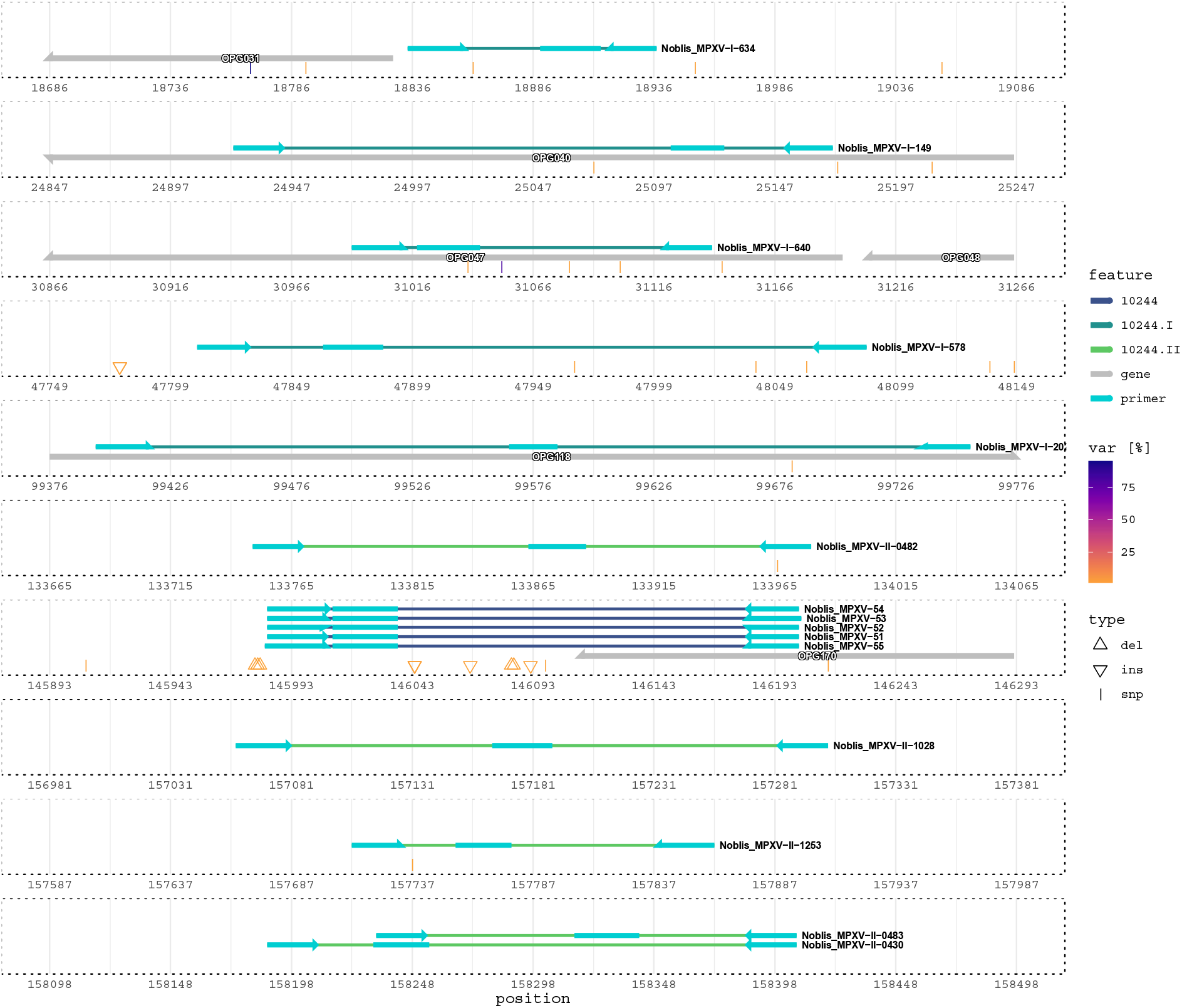
Genomic context of tested assays with respect to NCBI RefSeq accession NC_063383.1. Amplicon color corresponds to clade (10244 for generic mpox and a I or II suffix for each clade). Arrows and rectangles on each amplicon indicate primer and probe regions. Variations (indels and SNPs) are reported as a percentage of the 4,652 mpox sequences queried against the reference using nucmer.

**Table 1.**
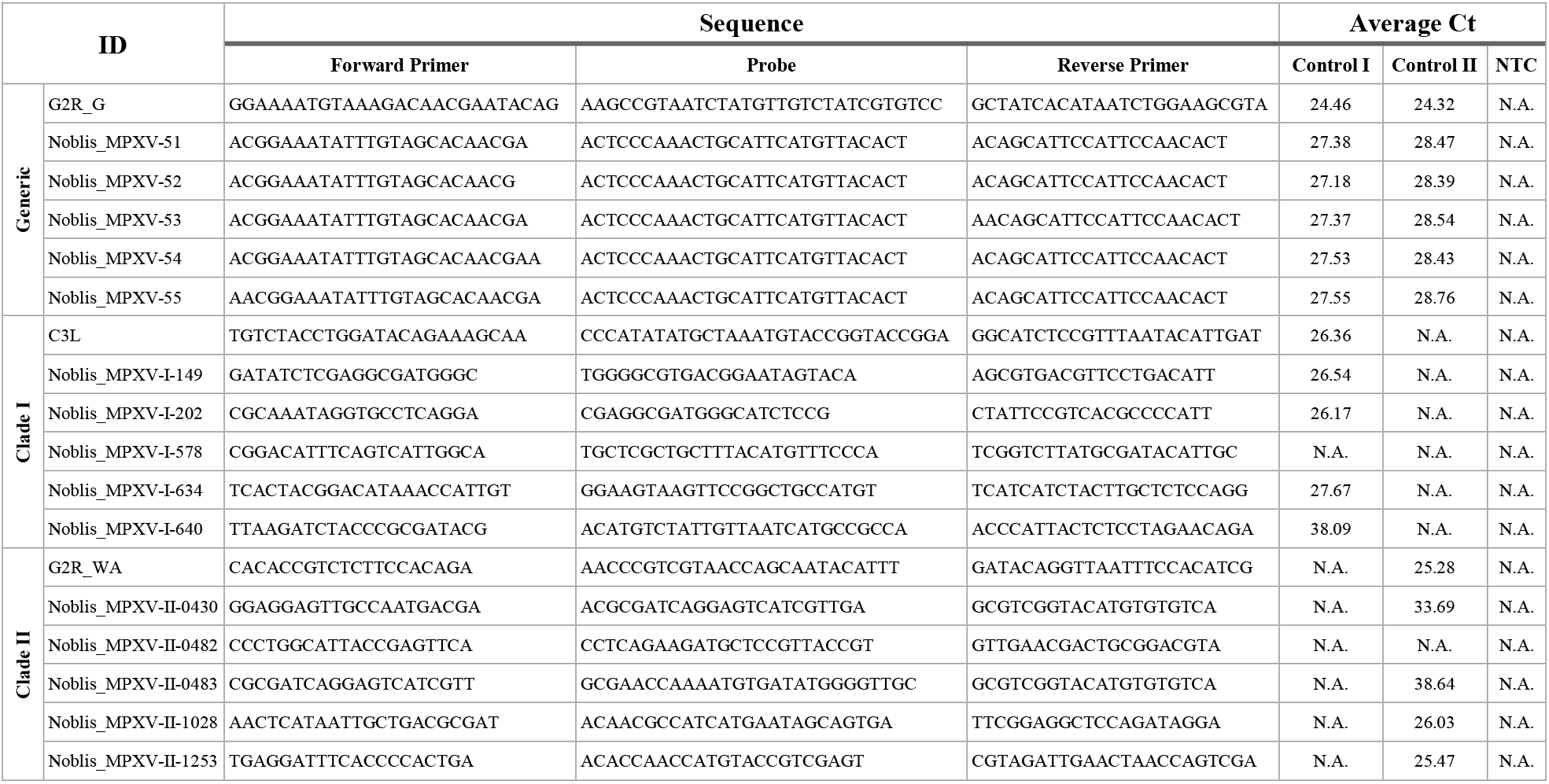
Control and Noblis, Inc. PCR assay primer/probe set sequences and average cycle threshold (Ct) values for control and no template control (NTC). Control I and Control II are controls for Clade I and Clade II. N.A. means no amplification. A copy of this table is available in Table S6.

Table 1 also lists the average cycle threshold (Ct) values and Figure 2 plots the normalized reporter values (Rn) for the published and candidate assays. The five generic candidate assays amplified both controls, but with less efficiency than the published assay. Two notably weak assays were observed with Ct values exceeding 38: Noblis_MPXV-I-640 and Noblis_MPXV-II-0483. All tested assays amplified their respective control targets except for Noblis_MPXV-I-578 (Clade I) and Noblis_MPXV-II-0482 (Clade II). The LOD for each assay tested was 5 copies/μL except for Noblis_MPXV-52 for Clade II, which was 50 copies/μL (Table S4 & Table S5). Figure S2 plots Rn for the LOD analysis.

**Figure 2.**
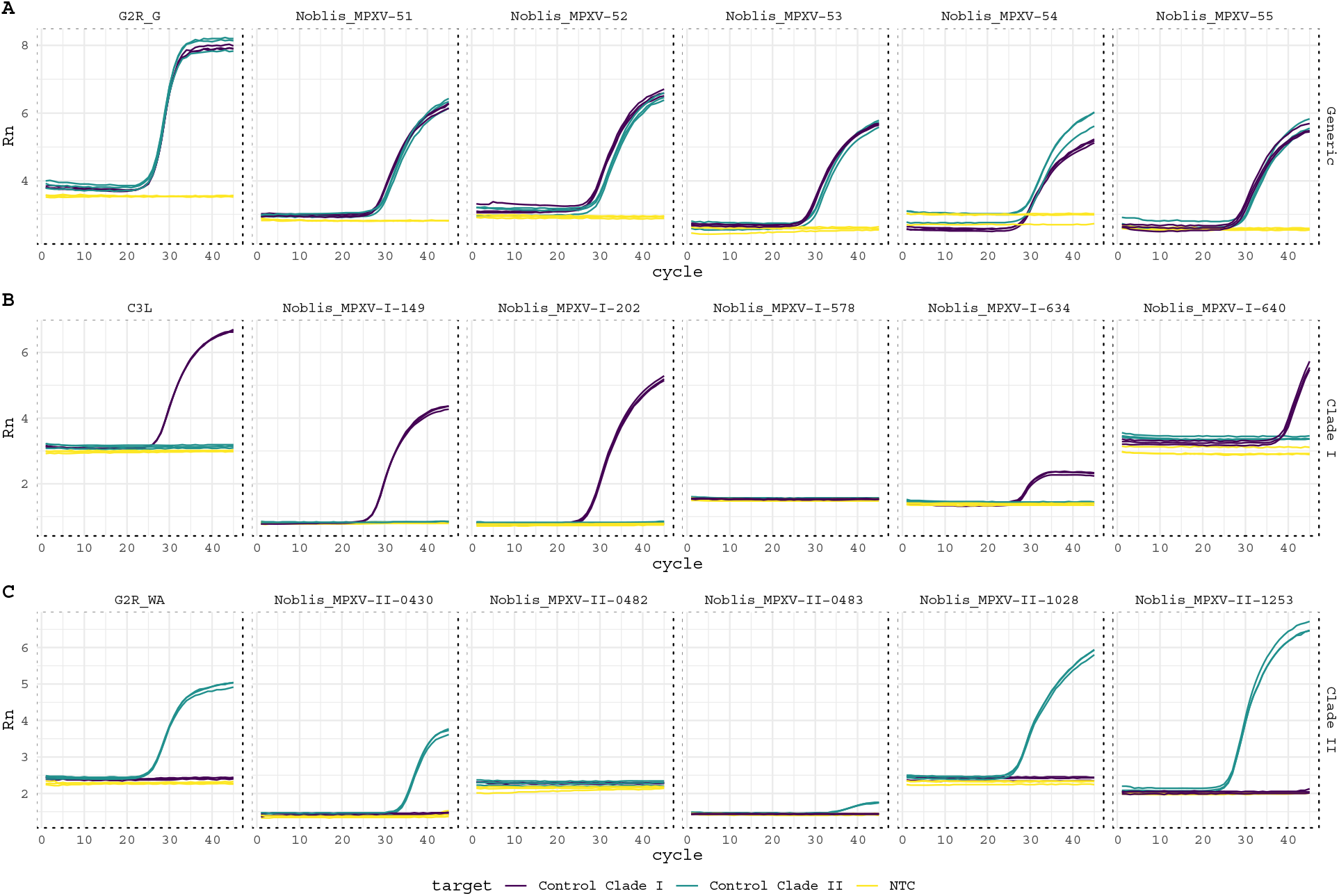
Amplification plots. Subplot rows indicate the intended target of each assay, with A, B, and C corresponding to generic pan-clade, Clade I, and Clade II.

## Discussion

This mpox study is the first to assess BioLaboro candidates *in vitro*. The objective was to demonstrate an enhanced ability to design PCR assays below the species level and to therefore identify clade-specific groupings. Previous work using the BioLaboro pipeline focused primarily on the PSET component to track predicted PCR assay performance over time and debug associated design issues. For example, a previous SARS-CoV-2 study demonstrated the sudden increase in failure rate of several assays related to deletions in the S gene as the virus evolved (17). This motivated the development of a new feature for PSET, which was the inclusion of 5’/3’-sequence context in the primer sequence query to increase local alignment quality and detect additional, potentially deleterious mutations outside of the binding target.

The mpox primer sets from the literature and those developed through this study were able to reliably detect *mpox virus*, except for Noblis_MPXV-I-578 and Noblis_MPXV-II-0483. The failure of the former is likely due to the false negative rate of 60.91% reported by PSET. This indicates that alignments failed immediately *in silico* at the local alignment step. Those remaining 43 alignments that passed the global-local threshold exhibited perfect identity except for one with an indel on the reverse primer. Noblis_MPXV-II-0483 had a near perfect true positive rate of 98.90% with the majority of the 4,047 alignments to the target clade having one substitution on the reverse primer. Remaining alignments exhibited several substitutions and indels, predominantly on the reverse primer. It is likely that the alignment threshold may have resulted in the calling of alignments with potentially deleterious mutations as true positives. Nevertheless, the overall success is likely a result of using strict BioVelocity and PSET thresholds.

*In vitro* tests showed that all assays amplified their respective clade targets, except for Noblis_MPXV-I-578 (Clade I) and Noblis_MPXV-II-0482 (Clade II). Overall, there were at least two Noblis generated primer sets per assay that produced comparable results to the published CDC assays. Optimization of Noblis assays has yet to be performed and could lead to better sensitivity than following the published protocols specific to the CDC assays. Outside of the Noblis_MPXV-52 Clade II assay, LOD testing of the best candidates from the original experiments were consistent with the results from the published assays. More testing is needed to demonstrate reproducibility and, therefore, improved confidence of the LOD data. Developing a multiplex PCR assay incorporating the optimal Generic, Clade I, and Clade II primer sets would be beneficial in advancing detection capability while providing a more efficient diagnosis option if made available to countries in need of this service.

As demonstrated in previous studies, clade-specific detection and resistance to false negatives due to primer sequence drift are critical to understanding the dimensions of an outbreak and coordinating a robust and effective public health response (17). The combination of algorithms in BioLaboro creates specific primer candidates and reliably assesses their performance *in silico* prior to in vitro testing which significantly contributes to a rapid diagnostic and public health response. Future studies should include testing on clinical samples and evaluation of assays in the field.

Our previous work focused primarily on the development of algorithms to calculate conserved DNA regions, generate candidate PCR primers, and monitor predicted PCR assay performance. This work demonstrates an end-to-end test of a workflow for the *in silico* generation and *in vitro* validation of novel assays. As proof-of-concept, we focused on developing generic and clade-specific *mpox virus* assays. The workflow was able to quickly provide viable assays with efficiencies comparable to those currently deployed. The rapid deployment of novel validated PCR assays benefits biosurveillance and pandemic response programs.

## Methods

### Pipeline

Noblis BioLaboro is a three-stage pipeline consisting of the BioVelocity, Primer3, and PSET (PCR Signature Erosion Tool) programs for the design and evaluation of PCR assays against reference DNA sequence data (18). BioVelocity calculates conserved genomic regions for a given set of sequence accessions or taxonomic identifiers. The algorithm builds a perfect-hash data structure mapping k-mer sequences to accessions. Conserved, overlapping k-mers unique to the target accessions are then extended with a mismatch threshold and combined into template sequences for use with Primer3 (2.6.1), which generates candidate PCR assays based on positional constraints, thermodynamic models, and secondary structure stability (19). Next, PSET further evaluates the assays via two-phased alignment (https://github.com/biolaboro/PSET). First, the workflow queries the amplicon against subjects using the blastn program of the BLAST+ suite (2.12.0+) and filters them with a combined percent identity and coverage threshold (20). Second, a global-local alignment step runs the glsearch36 program of the FASTA suite (36.3.8e) to check primer identity and orientation against the filtered set of subjects (21). The workflow outputs a confusion matrix based on alignment statistics and subject taxonomy. A true positive (TP) occurs when all assay primers align above the threshold with the correct arrangement on a subject bearing the targeted taxonomic identifier or descendant. If the taxonomy is incorrect, then a false positive (FP) is called. A true negative (TN) occurs when an assay fails alignment to a non-targeted subject and is a false negative (FN) otherwise. Note that sometimes these are estimates since the number of hittable subjects in the database is unknown due to the inconsistent usage of NCBI Taxonomy identifiers and varying levels of completeness and lengths of subject accessions.

### *In silico* Protocol

BioVelocity was run against the Bacterial (BCT) and Viral (VRL) database divisions of NCBI GenBank to map 50-mers and determine regions conserved within the mpox species (those bearing NCBI Taxonomy identifier 10244). Clade-specific regions were also targeted by selecting corresponding accessions provided by the Nextstrain monkeypox build (22). Primer3 was run using the conserved sequence output from BioVelocity and the configuration listed in Table S7. Prior to running PSET, an additional 10 nucleotides of 5’/3’-sequence context from a reference accession was added to the amplicon queries to enhance local alignment near the ends. PSET queried the context-padded amplicons using blastn against the NCBI nucleotide (nt) BLAST+ database and another built from exported GISAID EpiPox™ sequences (accession ranges listed in Table S8) (23). Assay primers were realigned using glsearch36 against the subject hits with ≥85% sequence similarity. Subsequent hits to subjects with ≥90% sequence similarity and correct taxonomy were considered TPs. PSET also evaluated assays from the literature, including one pan-orthopox (E9L), two mpox assays (B6R and G2R_G), one Clade I (C3L), and one Clade II (G2R_WA) with the same procedure (24, 25) (Table S1).

Confusion matrix values were calculated by taxon (species) and clade based on results from NCBI nt and GISAID EpiPox™. For the former, no assumptions were made about the total number of potential positive hits. This was done for the latter since only high-quality, complete sequences were downloaded. Accordingly, there were 110 and 4,454 potential positives for Clade I and Clade II and 4,652 for generic assays (including those subjects with unknown clade). Note that the FP rate (FPR) and TN rate (TNR) for generic assays are undefined with respect to the GISAID EpiPox™ database since no negative hits are possible. Candidate assays for testing were selected based on Primer3 penalty point values, confusion matrix results, and alignments.

Genomic variation was calculated using the nucmer program of the MUMMER4 suite to determine single-nucleotide polymorphisms (SNPs) and insertions/deletions (indels) (26). NCBI RefSeq accession number NC_063383.1 served as the reference and the exported GISAID EpiPox™ sequences comprised the query (Table S8). The program was run with default parameters.

### *In vitro* Protocol

Previously published primer sets were run alongside Noblis, Inc. candidates to confirm successful real-time PCR tests, including G2R_G (generic), C3L (Clade I), and G2R_WA (Clade II) (Table 1) (24, 25). Synthetic human monkeypox (hMPXV) controls from Twist Bioscience were used for the validation of the Noblis assay candidates, with Control 1 (Part No. 106056) and Control 2 (Part No. 106059) specific to Clade I and Clade II strains respectively. Stock concentration for each control was 10^5^ copies/μL. Dilutions of 1:20 were performed (5 μL control into 95 μL molecular grade water) to give a final concentration of 5·10^3^ copies/μL. The fifteen final candidates and published sets for each assay were ordered as PrimeTime Std qPCR Assays through Integrated DNA Technologies (IDT) and used with the PrimeTime Gene Expression Master Mix. PrimeTime Std qPCR Assays were resuspended in 500 μL TE for a 20X stock concentration. 200 μL of reference dye was added to the PrimeTime Gene Expression 2X Master Mix per IDT’s instruction for use with the Applied Biosystems StepOnePlus thermocycler. Master mix setup for each reaction included 10 μL of the PrimeTime Gene Expression 2X Master Mix, 1 μL of the appropriate PrimeTime Std qPCR Assay and 4 μL of molecular grade water. 96-well plates were used with the 15 μL master mix added to the necessary wells. 5 μL of the appropriate template was then added to the wells and all samples were tested in triplicate. Conditions were 1 cycle at 95°C for 3 min, 45 cycles at 95°C for 5 sec, and 60°C (62°C for Clade II) for 30 sec. Limit of detection (LOD) was determined for the best assays based on average Ct from the original tests using the aforementioned parameters. Ten-fold serial dilutions were created from the 1:20 initial dilution of each control out to 1:2,000,000.

## Data Availability

This study relied on genomic DNA sequence data and metadata available at NCBI GenBank (https://www.ncbi.nlm.nih.gov/genbank/), GISAID EpiPox™ (https://gisaid.org/), and Nextstrain (https://nextstrain.org/monkeypox/hmpxv1). Table S8 lists the GISAID EpiPox™ accession numbers. Code, sequence, and raw PCR data are also available on Figshare (10.6084/m9.figshare.23800629). Please contact the corresponding author with requests for additional information.

## Author Contributions

Conceptualization: DN, NT, SG, CP, BA, & DA; Data Curation: DN, NT, & SM; Formal analysis: DN, NT, & SM; Funding acquisition: KJ & DA; Investigation: DN, NT, & SM; Methodology: DN, NT, CP, BA, & DA; Project administration: DN, NT, KJ, & DA; Resources: SG, SWR, JH, KJ, BA, & DA; Software: DN, SM, & BA; Supervision: KJ & DA; Validation: DN, NT, SG, & DA; Visualization: DN, NT, SG, & SWR; Writing - Original Draft: DN, NT, SWR, & BA; Writing - Review & Editing: DN, NT, SG, SWR, CP, KJ, BA, LB, & DA. Contributions and contact information are listed in Table SA and Table SB.

## Acknowledgements

We gratefully acknowledge all data contributors, i.e., the Authors and their Originating laboratories responsible for obtaining the specimens, and their Submitting laboratories for generating the genetic sequence and metadata and sharing via the GISAID Initiative, on which this research is based. Noblis, Inc. funded this study as part of the internal Noblis Sponsored Research (NSR) program. Many thanks to Dr. Shanmuga Sozhamannan (ORCiD:0000-0002-6762-7874) for feedback on the PSET algorithm and its development. We also thank Dr. Michael Wiley (ORCiD:0000-0001-6688-007X) for feedback on the results and ideas for continued development and deployment.

---

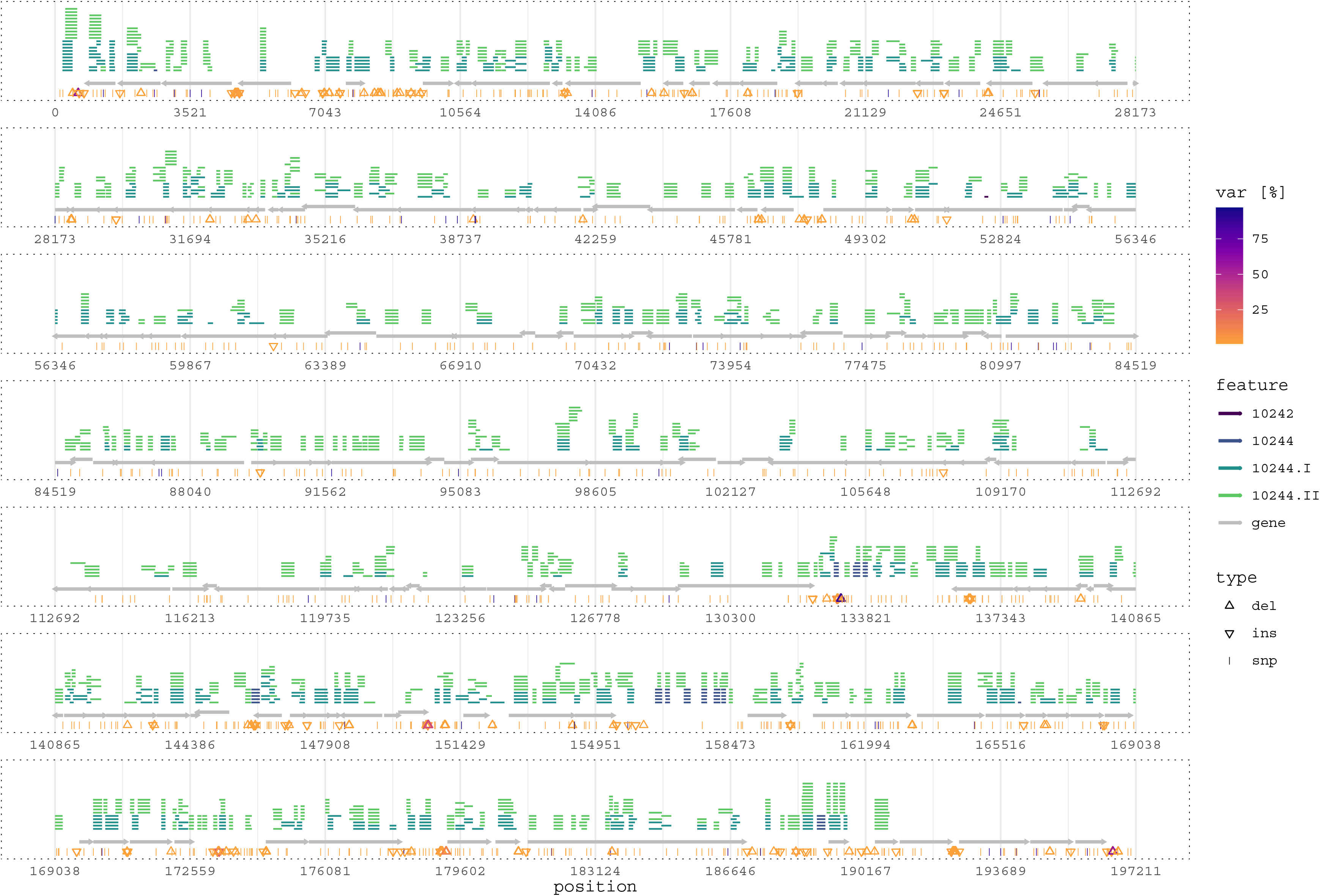

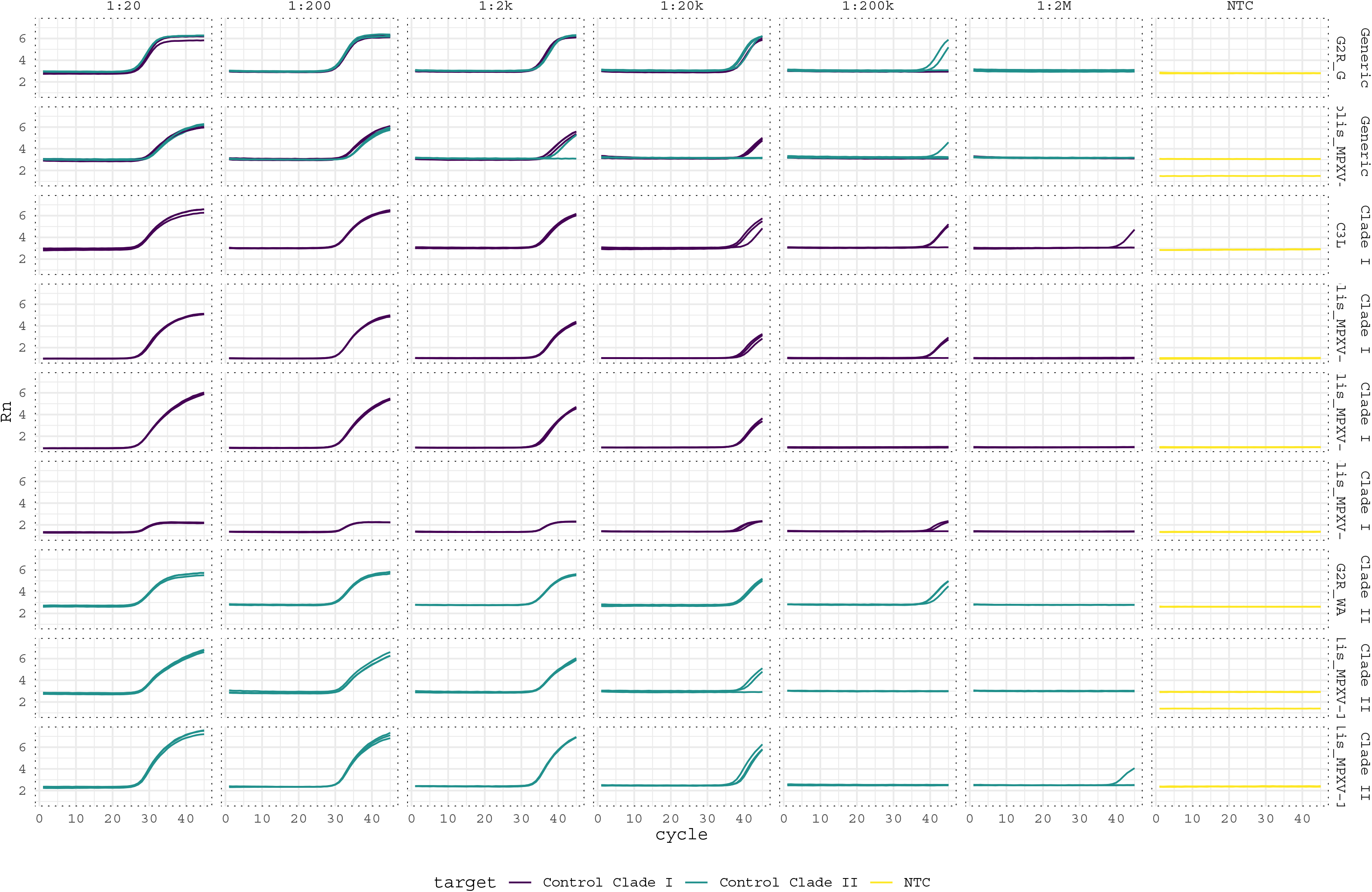

## Notes

https://www.ncbi.nlm.nih.gov/genbank

https://gisaid.org/

https://nextstrain.org/monkeypox/hmpxv1

https://figshare.com/articles/software/Clade-Specific_MPXV_PCR_Assays_Code_and_Materials_Supplement/23800629

https://github.com/biolaboro/pset

### Competing Interest Statement

The authors have declared no competing interest.

### Summary of Updates

- Adding limit of detection results - Adding importance section to the abstract - Formatting for submission to mSphere - Including additional supplementary materials - Add data availability statement

